# Caspase-4/11 exacerbates disease severity in SARS-CoV-2 infection by promoting inflammation and thrombosis

**DOI:** 10.1101/2021.09.24.461743

**Authors:** Mostafa Eltobgy, Ashley Zani, Adam D. Kenney, Shady Estfanous, Eunsoo Kim, Asmaa Badr, Cierra Carafice, Kylene Daily, Owen Whitham, Maciej Pietrzak, Amy Webb, Jeffrey Kawahara, Adrian C. Eddy, Parker Denz, Mijia Lu, KC Mahesh, Mark E. Peeples, Jianrong Li, Jian Zhu, Jianwen Que, Richard Robinson, Oscar Rosas Mejia, Rachael E. Rayner, Luanne Hall-Stoodley, Stephanie Seveau, Mikhail A. Gavrilin, Andrea Tedeschi, Santiago Partida-Sanchez, Frank Roberto, Emily A. Hemann, Eman Abdelrazik, Adriana Forero, Shahid M. Nimjee, Prosper Boyaka, Estelle Cormet-Boyaka, Jacob S. Yount, Amal O. Amer

## Abstract

SARS-CoV-2 is a worldwide health concern, and new treatment strategies are needed ^1^. Targeting inflammatory innate immunity pathways holds therapeutic promise, but effective molecular targets remain elusive. Here, we show that human caspase-4 (CASP4), and its mouse homologue, caspase-11 (CASP11), are upregulated in SARS-CoV-2 infections, and that *CASP4* expression correlates with severity of SARS-CoV-2 infection in humans. SARS-CoV-2-infected *Casp11*^-/-^ mice were protected from severe weight loss and lung pathology, including blood vessel damage, compared to wild-type (WT) and gasdermin-D knock out (*Gsdmd*^*-/-*^) mice. GSDMD is a downstream effector of CASP11 and CASP1. Notably, viral titers were similar in the three genotypes. Global transcriptomics of SARS-CoV-2-infected WT, *Casp11*^-/-^ and *Gsdmd*^*-/-*^ lungs identified restrained expression of inflammatory molecules and altered neutrophil gene signatures in *Casp11*^-/-^ mice. We confirmed that protein levels of inflammatory mediators IL-1β, IL6, and CXCL1, and neutrophil functions, were reduced in *Casp11*^-/-^ lungs. Additionally, *Casp11*^-/-^ lungs accumulated less von Willebrand factor, a marker for endothelial damage, but expressed more Kruppel-Like Factor 2, a transcription factor that maintains vascular integrity. Overall, our results demonstrate that CASP4/11, promotes detrimental SARS-CoV-2-associated inflammation and coagulopathy, largely independently of GSDMD, identifying CASP4/11 as a promising drug target for treatment and prevention of severe COVID-19.

## Main

Severe acute respiratory syndrome coronavirus 2 (SARS-CoV-2) is the causative infectious agent of the worldwide COVID-19 pandemic^1^. SARS-CoV-2 is a positive sense single-stranded RNA virus that can induce hyper-inflammatory responses, including cytokine storm, in the lungs as well as extra-pulmonary organs in severe cases^2^. IL-6, CXCL1, IL-1α, IL-1β and type I interferons, among other cytokines, are thought to contribute to pathological manifestations of the SARS-CoV-2 infection^3^. In addition, formation of thrombi that can cause myocardial infarction, stroke and pulmonary embolism is a hallmark of severe Covid-19. Endothelial and neutrophil dysfunctions during SARS-CoV-2 infection increase the incidence of thromboembolic complications^4^. Thrombus formation is initiated by von Willebrand factor (VWF), a glycoprotein released by damaged endothelial cells and megakaryocytes^5,6^. VWF also self-associates, forming strings protruding into the lumen serving as a scaffold for platelet adhesion and aggregation^6^. Cellular sensors of infection, such as Toll-like receptor 2 (TLR2), C-type lectin receptors, and the NLRP3 inflammasome have been implicated in triggering the induction and secretion of cytokines and inflammatory lung damage in SARS-CoV-2 infections^7^. However, the contribution of these pathogen-sensing pathways and other inflammasome components in mediating host defense versus immune-mediated pathology and thrombosis during SARS-CoV-2 infection *in vivo* remains unclear^7^. While effector molecules downstream of infection-sensing pathways, such as specific inflammatory cytokines, have been targeted in attempts to limit virus-induced tissue damage, most of these strategies failed to exert major benefits in human clinical trials^8^. Therefore, strategies targeting molecules upstream of multiple inflammatory cytokines or chemokines may be more effective, though this remains to be experimentally tested. Here, we investigate the role of a major member of the non-canonical inflammasome, caspase-11 (CASP11), and its downstream effector Gasdermin D (GSDMD) in SARS-CoV-2 infection and disease severity using knockout mouse models and mouse-adapted SARS-CoV-2.

Caspases are a family of cysteine proteases that specifically cleave their substrates at the C-terminal side of aspartic acid residues. CASP11 is a murine protein that is critical for defense against bacterial pathogens. Human caspase-4 (CASP4) displays high homology to murine CASP11^9,10^ and we have demonstrated that human CASP4 mediates many functions of mouse CASP11 in macrophages during bacterial infections^9^. CASP4/11 is a component of the non-canonical inflammasome with multiple functions that remain to be fully characterized. One major role for this protein is the cleavage of GSDMD^11^. Once cleaved, the GSDMD N-terminal fragment inserts into the plasma membrane of eukaryotic cells to form pores that allow the release of IL-1β and other molecules, sometimes leading to cell lysis and death known as pyroptosis^12^. Interestingly, accumulating evidence has posited potential roles for GSDMD downstream of caspases in mediating inflammatory pathology during SARS-CoV-2 infection^13^. SARS-CoV-2 infection studies in GSDMD genetically deficient animal models have not yet been performed, though clinical trials testing inhibitors of GSDMD in COVID-19 patients were not promising^8^. Likewise, the role of CASP4/11 in viral infections has not been explored, despite the induction of these proteins by the antiviral type I and II interferons^8^. This notable induction by interferons and the broad roles of CASP4/11 in regulating diverse inflammatory pathologies, including bacterial infections, gouty arthritis, and gastroenteritis^10^, prompted us to investigate its role in SARS-CoV-2 infections.

## Results

### CASP4/11 expression is elevated in the lungs during SARS-CoV-2 infections of mice and humans and correlates with disease severity in humans

CASP4/11 is weakly expressed or absent in resting cells, but is induced in response to bacterial infections^14^. The analysis of publicly available RNA sequencing data of nasopharyngeal swab material from subjects with SARS-CoV-2 and healthy donors (GEO accession: GSE163151), revealed that *CASP4* is highly expressed in the airway of SARS-CoV-2-infected patients, and that expression levels increase with disease severity (**Fig. 1a**). Additionally, we found that human lung sections from COVID-19 patients show higher levels of CASP4 staining compared with healthy lung controls (**Fig. 1b**), owing to greater numbers of CASP4 positive cells in the infected lung tissue (**Fig. 1c**). We then performed intranasal infection of C57BL/6 wild-type (WT) mice with pathogenic mouse-adapted SARS-CoV-2 (strain MA10)^15^, and found that infection strongly induces *Casp11* expression throughout murine lung tissue within 4 days of infection as detected by RNAscope *in situ* hybridization (ISH) (**Fig. 1d**) and confirmed by qRT-PCR (**Fig. 1e**). The level of CASP11 protein, likewise, went from below detection to highly-expressed in response to SARS-CoV-2 infection of murine lungs (**Fig. 1f**). We further examined infection of K18-hACE2 mice expressing the human ACE2 receptor using human isolate SARS-CoV-2 strain USA-WA1/2020 (WA1). Similar to mouse adapted SARS-CoV-2, the non-adapted human virus strongly induced the lung expression of CASP11 as demonstrated by qRT-PCR (**Fig. 1g**)^16^. Overall, CASP4 is highly expressed in the lungs of COVID-19 patients, and CASP11 is similarly induced upon SARS-CoV-2 infection of mice.

**Figure 1:**
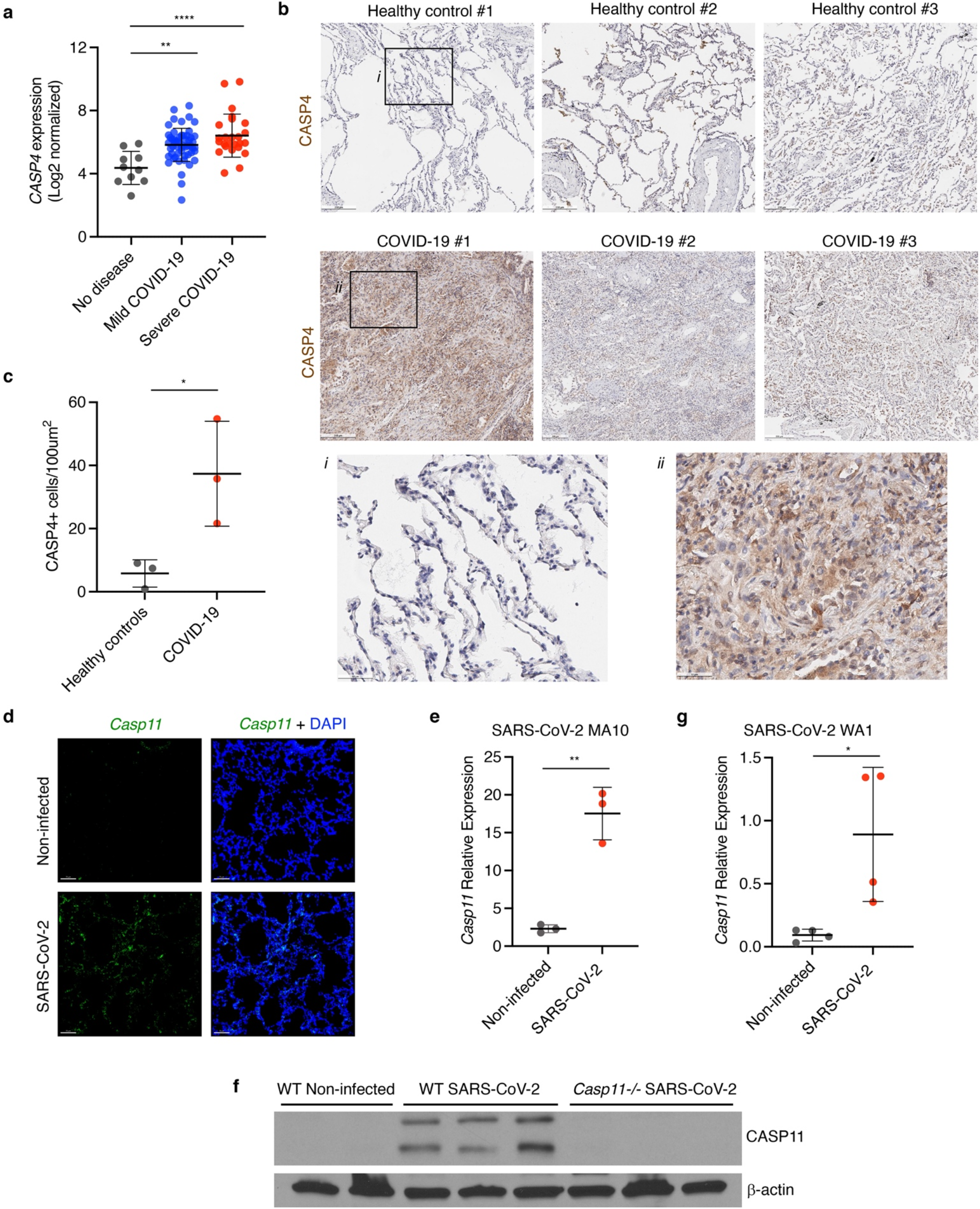
CASP4 is upregulated in humans and mice infected with SARS-CoV-2. **a**, *CASP4* expression levels from RNA sequencing of nasopharyngeal swab samples from patients with no disease, mild SARS-CoV-2, or severe SARS-CoV-2 [GSE145926], one way ANOVA with Tukey’s multiple comparisons test. **b**, Human lung samples from 3 donors with healthy lungs or from 3 donors who died of SARS-CoV-2 were stained for CASP4 (brown). Black boxes (*i, ii*) outline zoomed regions. **c**, Quantification of CASP4 positive cells from lungs in **b**, unpaired t test. **d-f**, Mice were infected for 4 days with mouse adapted SARS-CoV-2 (MA10, 10^5^ pfu). **d**, *Casp11* RNA (green, RNAscope *in situ* hybridization) and DAPI (blue) were visualized (3D Intensity projection image) in lung sections using 20x objective. **e**, *Casp11* RNA levels were quantfied in lung samples (N=3) by qRT-PCR, unpaired t test. **f**, CASP11 protein levels in lungs described in **d** (N=3) were examined by Western blot. **g**, K18-hACE2 mice were infected for 4 days with human SARS-CoV-2 (WA1, 10^5^ pfu) and *Casp11* RNA levels were quantitated in lung samples (N=4) by qRT-PCR, unpaired t test. *p<0.05, **p<0.005, ****p<0.0001.

### *Casp11* deficiency reduces disease severity in SARS-CoV-2-infected mice

We next examined whether CASP11 regulates disease severity caused by SARS-CoV-2 infection. Wild-type (WT), *Casp11*^-/-^ and *Gsdmd*^-/-^, and mice were infected with SARS-CoV-2 MA10 for comparison of weight loss, a commonly used indicator of overall infection severity in mice^16^. We found that WT mice lost a significant percentage of their body weight between days 1 and 4 post-infection, followed by partial recovery of weight up to day 7, at which point we ended our experiments (**Fig. 2a**). *Casp11*^-/-^ mice, on the other hand, lost weight only up to day 3, and then rapidly recovered fully to their original weight by day 5 (**Fig. 2a**). In comparison, weight loss of *Gsdmd*^-/-^ mice was not significantly different from that of WT mice (**Fig. 2a**). These data indicate that CASP11 promotes disease severity during SARS-CoV-2 infection, and that this function is not mediated by GSDMD.

**Figure 2:**
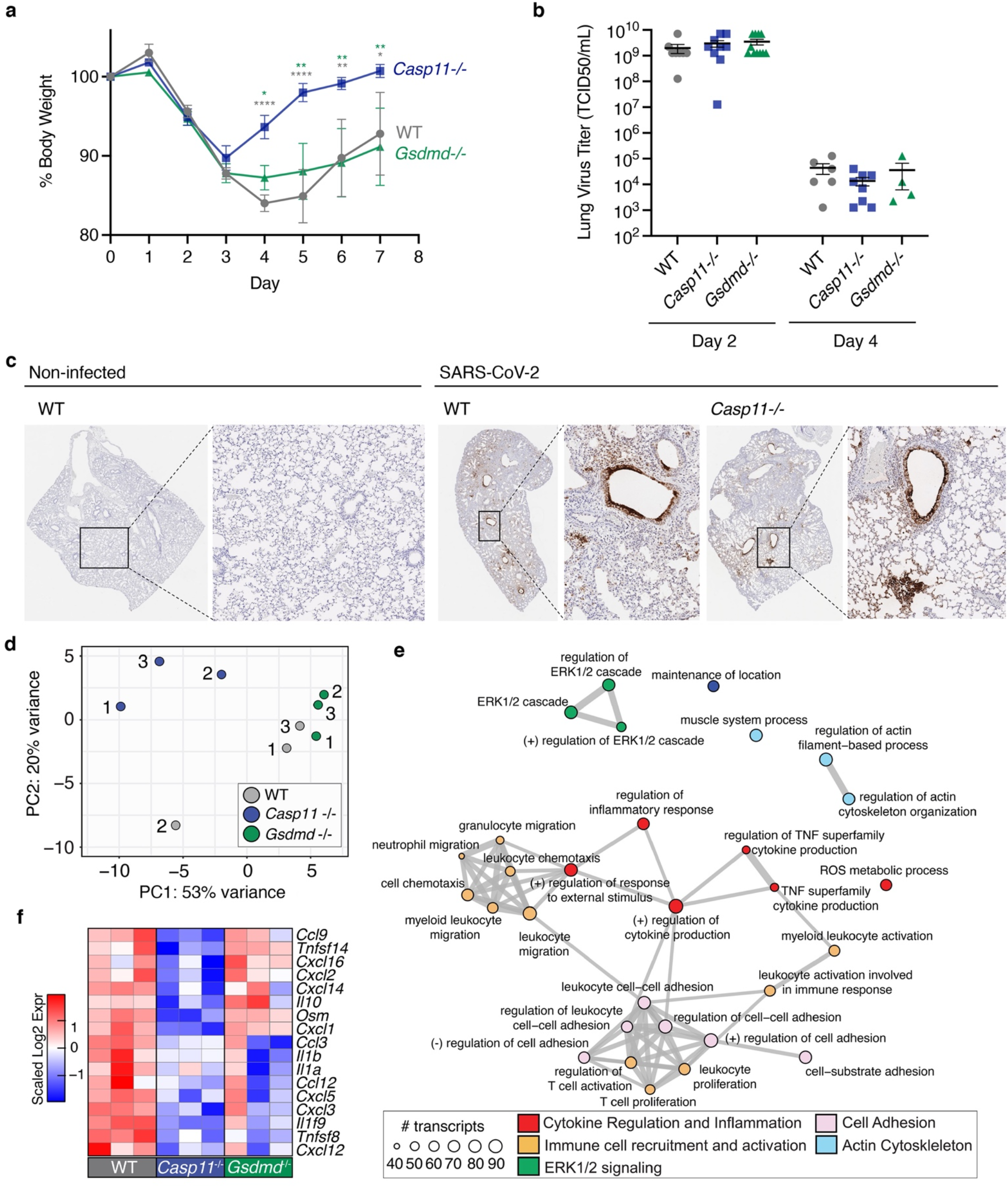
*Casp11*^-/-^ mice show decreased SARS-CoV-2 infection severity without affecting viral titers but by modulating specific inflammatory programs. **a-c**, WT, *Casp11*^*-/-*^ and *Gsdmd*^*-/-*^ mice were infected with SARS-CoV-2 (MA10, 10^5^ pfu). Weight loss was tracked for 7 d, *p<0.05, **p<0.005, ****p<0.0001, ANOVA with Bonferonni’s multiple comparisons test (**a**), Day 0-4 WT (N=7), *Casp11*^*-/-*^ (N=10), *Gsdmd*^*-/-*^ (N=9); Day 5-7 WT (N=4), *Casp11*^*-/-*^ (N=7), *Gsdmd*^*-/-*^ (N=6). **b**, TCID50 viral titers were quantified in lung tissue homogenates. **c**, Sections from non-infected control lungs or lungs collected at 4 days post-infection were stained for SARS-CoV-2 nucleocapsid protein (brown staining, images representative of at least 3 mice per group). **d-f**, WT, *Casp11*^*-/-*^, and *Gsdmd*^*-/-*^ mice (N=3) were infected with SARS-CoV-2 (MA10, 10^5^ pfu) for 2 days. RNA was extracted from lungs and subjected to RNA sequencing. **d**, Principal component analysis (PCA) of SARS-CoV2-infected lung gene expression with points representing individual WT (grey), *Casp11*^-/-^(blue), and *Gsdmd*^-/-^ (green) mice. **e**, Top 30 significant Gene Ontology Biological Pathways are depicted. Node size indicates the number of transcripts within each functional category. Edges connect overlapping gene sets. Numbers represent individual replicates and color indicates relative upregulation (red) or downregulation (blue) in gene expression. **f**, Heatmap of significantly changed cytokine and chemokine genes when comparing *Casp11*^-/-^ infected lungs versus WT. Expression scaling is relative to WT and *Gsdmd*^*-/-*^ mice for comparisons (N=3) (p<0.05)..

To determine whether differences in disease severity could be explained by differences in viral replication, we quantified live virus titers in WT, *Casp11*^-/-^ and *Gsdmd*^-/-^ mouse lungs at 2 and 4 days post-infection. We found that viral loads were similar with no statistical difference between the mice genotypes at either time point (**Fig. 2b**). We also observed that, in agreement with previous reports^17^, viral titers were decreased at day 4 compared with day 2 in all groups, demonstrating that neither CASP11 nor GSDMD are required for viral clearance mechanisms (**Fig. 2b**). To corroborate these findings, lung sections from WT and *Casp11*^-/-^ mice were stained for SARS-CoV-2 nucleocapsid protein and similar staining patterns were observed with prominent infection of cells lining the airways and neighboring alveoli (**Fig. 2c**). Overall, these results demonstrate that loss of CASP11, but not GSDMD, prevents severe disease in SARS-CoV-2 infection without affecting virus replication or clearance.

### *Casp11*^*-/-*^ SARS-CoV-2-infected lungs elicit specific inflammatory gene signatures

To examine global transcriptional effects of CASP11 and GSDMD in the lung during SARS-CoV-2 infections, we infected WT, *Casp11*^-/-^ and *Gsdmd*^-/-^ mice, and performed RNA sequencing on lung RNA at 2 days post-infection. Day 2 was chosen because it is the peak of virus replication in the lungs of mice^15^. No similarity between the gene signatures obtained from WT, *Casp11*^-/-^ and *Gsdmd*^-/-^ infected lungs were seen using dimensionality reduction approaches, while the *Casp11*^-/-^ infected lung profiles showed greater divergence in gene expression patterns (**Fig. 2d**). We contrasted the significant gene expression changes (p-value <0.05) in infected *Casp11*^-/-^ and *Gsdmd*^-/-^ lungs relative to WT mice to understand how these deficiencies impact the transcriptional landscape in terms of differentially expressed genes (DE, either significantly upregulated or downregulated) (**Supplementary Fig. 1a**). Functional analysis of DE genes in *Casp11*^-/-^ versus WT lungs revealed an enrichment for genes corresponding to immunological pathways involved in cytokine production and inflammation (red), immune cell migration and activation (orange), cell adhesion (pink), and ERK1/2 signaling (green) (**Fig. 2e**). In accordance with known actin polymerization regulation imparted by CASP11, the absence of *Casp11* in SARS-CoV-2 infection also resulted in changes in genes involved in actin regulatory pathways (blue) (**Fig. 2e**)^9,18,19^. Since sensing of virus replication by cells generally induces interferon (IFN)-mediated antiviral responses and the expression of inflammatory cytokines, we investigated whether CASP11 or GSDMD shape the antiviral gene program during SARS-CoV-2 infection. First, we specifically examined the expression of IFN-stimulated genes (ISGs), which are abundantly upregulated by type I IFN stimulation in murine airway epithelial cells^20^. Deficiency of CASP11 or GSDMD did not result in differential ISG (LFC |0.58|; p-value <0.05) expression relative to WT infected lungs (**Supplementary Fig. 1b,c**).

Specific examination of cytokine and chemokine genes revealed a statistically significant downregulation of several important inflammatory mediators in the absence of *Casp11* including cytokines *Il1b, Il1α*, and *Il1f9*, and chemokines *Cxcl1, Cxcl2, Cxcl14, Cxcl3, Cxcl5*, and *Ccl3* (**Fig. 2f**). These findings are consistent with ERK activation downstream of CXCL1 and CXCL3 signaling as highlighted in (**Fig. 2e)**. Knockout of *Gsdmd*, had less impact on the magnitude of cytokine and chemokine expression compared to *Casp11* knockout (**Fig. 2f**). Overall, our results demonstrate that CASP11 controls a specific subset of inflammatory responses during SARS-CoV-2 infection.

### CASP11 promotes the production of specific inflammatory mediators in response to SARS-CoV-2 *in vivo* and *in vitro*

To examine the role of CASP11 in mediating the pathological hallmarks of SARS-CoV-2 pulmonary infection, lung sections from infected WT, *Casp11*^-/-^ *and Gsdmd*^-/-^ mice were fixed and stained with hematoxylin and eosin (H&E). Sections from all infected animals showed areas of consolidated lung tissue indicative of cellular infiltration and inflammation that was absent in non-infected control tissue (**Fig 3a**). However, WT and *Gsdmd*^-/-^ lung sections showed more severe tissue consolidation and cell infiltration throughout a greater portion of the lung than that seen in *Casp11*^-/-^ mice. We thus quantified cell area versus airway space to determine cellularity scores indicative of pathology for tissue sections from individual mice^21^. We observed significantly decreased SARS-CoV-2-induced lung pathology in *Casp11*^-/-^ mice compared to WT and *Gsdmd*^*-/-*^ mice (**Fig. 3a,b**), correlating with the preservation of *Casp11*^-/-^ mice body weight and their faster recovery that (**Fig. 2a**).

**Figure 3:**
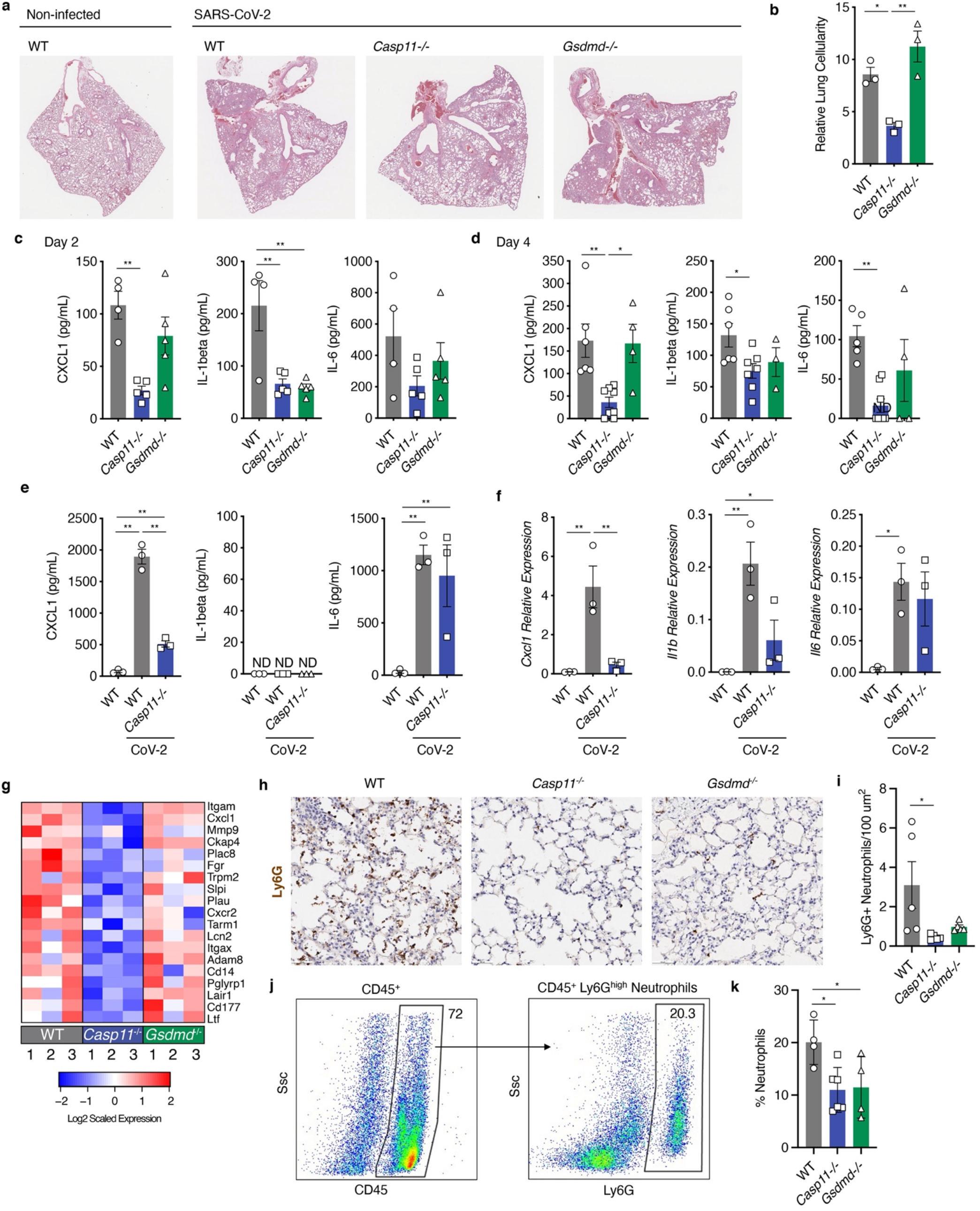
*Casp11*^*-/-*^ mice show decreased lung inflammation, less neutrophil recruitment, and altered neutrophil function in response to SARS-CoV-2 infection. **a-b**, WT and *Casp11*^*-/-*^ mice were infected with SARS-CoV-2 (MA10, 10^5^ pfu). **a**, Lung sections from day 4 post-infection were stained with H&E to visualize lung damage and airway consolidation. **b**, Lung sections as in **a** were analyzed by the color deconvolution method to quantify cellularity as an indicator of cellular infiltration and alveolar wall thickening, ANOVA with Tukey’s multiple comparisons test. **c,d**, Lung homogenates from 2 or 4 days post-infection were analyzed by ELISA for detection of CXCL1, IL-1β, or IL-6, ANOVA with Tukey’s multiple comparisons test. **e,f**, Macrophages were purified from lungs of mice of the indicated genotype. The cells were infected with mouse adapted SARS-CoV-2 (MOI 1) for 24 h. Cell supernatants were analyzed by ELISA, or cellular RNA was analyzed by qRT-PCR for the indicated chemokine/cytokines, ANOVA with Tukey’s multiple comparisons test. **g**, Heatmap of significantly changed neutrophil-related genes comparing *Casp11*^-/-^ infected lungs versus WT(p<0.05). Expression scaling is relative to WT and *Gsdmd*^*-/-*^ mice for comparisons. Numbers represent individual replicates and color indicates relative upregulation (red) or downregulation (blue) in gene expression. **h**, Lung sections of day 2 SARS-CoV-2-infected WT, *Casp11*^-/-^ and *Gsdmd*^*-/-*^mice (N=5) stained with neutrophil marker Ly6G and quantified in **i. j**, Flow cytometry of lungs homogenates from mice WT (N=4), Casp11^-/-^ (N=6), Gsdmd^-/-^ (N=4) as in **h** and quantified in **k**. *p<0.05, **p<0.005.

Guided by our transcriptomic results indicating that a critical subset of inflammatory mediators are controlled by CASP11 (**Fig. 2f**), we measured levels of CXCL1, IL-1β, and IL-6 by ELISA in lung homogenates from infected animals at 2 and 4 days post-infection (**Fig. 3c,d**). IL-1β was lower in the lungs of both *Casp11*^-/-^ and *Gsdmd*^-/-^ mice at 2 days post-infection when compared to WT (**Fig. 3c)**. Moreover, IL-1β staining in lung tissue sections revealed more IL-1β in WT SARS-CoV-2 infected mice than *Casp11*^-/-^ infected ones (**Supplementary Fig. 2**). On the other hand, the production of CXCL1 was dependent on *Casp11* at both time-points, and independent of *Gsdmd* (**Fig. 3c,d**). Average levels of IL-6 were partially decreased in *Casp11*^*-/-*^ lungs with a statistically significant difference between WT and *Casp11*^*-/-*^ lungs at day 4 (**Fig. 3c,d**). These results corroborate and expand our day 2 transcriptomic analysis in which expression of *Il1b* and *Cxcl1* was decreased (**Fig. 2f**), and demonstrate that production of a critical subset of inflammatory mediators in the lung is dependent on CASP11 during SARS-CoV-2 infection.

To determine the role of CASP11 in the response of lung macrophages to SARS-CoV-2, we purified mature primary macrophages from lungs of WT and *Casp11*^-/-^ mice, and infected them with SARS-CoV-2 MA10. Culture supernatants and cellular RNA were collected and measured for IL-1β, IL-6 and CXCL1 protein and transcript levels, respectively. Compared with non-infected cells, CXCL1 protein and RNA transcripts were detected at high levels upon infection of WT macrophages, but were poorly produced by *Casp11*^*-/-*^ cells (**Fig. 3e,f**). Interestingly, IL-1β transcripts were also induced in a CASP11-dependent manner, but secreted protein was not detected in either group (**Fig. 3e,f**). Distinctly, protein and transcript levels of IL-6 did not significantly differ between WT and *Casp11*^*-/-*^ cells (**Fig. 3e,f**). These results confirm our *in vivo* measurements and further demonstrate that CASP11 is an important cellular regulator of specific cytokines and chemokines, including CXCL1 and IL-1β, in response to SARS-CoV-2.

### CASP11 promotes lung neutrophil responses during SARS-CoV-2 infection

To better understand the biological processes regulated by *Casp11*, we further analyzed the functional gene enrichment categories of the 236 genes most downregulated (LFC <-0.58; p-value <0.05) in *Casp11*^*-/-*^ lungs. A striking enrichment of neutrophil-related gene signatures emerged that included neutrophil-specific markers (e.g., *Cd177* and *Cxcr2*), neutrophil degranulation genes (e.g., *Pglryp1, Ckap4, Adam8*, and *Plac8*), and neutrophil complement receptors (*Itgam* and *Itgax*), among others (**Fig. 3g, Supplementary Fig. 1d**). Additionally, genes associated with the response to tissue damage from neutrophils (*Slpi* and *Lair1*) were also decreased in the absence of *Casp11* relative to WT lungs (**Fig. 3g**). These results are consistent with decreased gene expression for the neutrophil chemoattractant CXCL1 (**Fig. 3c,d**), as well as with previous reports of neutrophil regulation by CASP11 through effects on actin^9,19^.

Notably, expression of these neutrophil signature genes in *Gsdmd*^*-/-*^ lungs was less affected than in *Casp11*^*-/-*^ lungs (**Fig. 3g)**, though other genes that are downregulated in the absence of *Gsdmd* were weakly associated with dysregulation of other immune pathways (**Supplementary Fig. 1e**). Conversely, analysis of genes upregulated in the absence of *Casp11* revealed a putative association with muscle-specific pathways (**Supplementary Fig. 1f**), while genes most upregulated in *Gsdmd*^*-/-*^ lungs were not enriched for any specific functional pathways. Overall, these analyses most prominently demonstrate that CASP11 is required for robust production of specific inflammatory mediators as well as neutrophil recruitment and functions in the lung during SARS-CoV-2 infection.

To further examine the role of neutrophils in SARS-CoV-2 infection, lung sections from WT, *Casp11*^*-/-*^ and *GsdmD*^*-/-*^ were stained for the neutrophil marker Ly6G (**Fig. 3h**). Quantification of Ly6G positive cells demonstrated fewer neutrophils in *Casp11*^*-/-*^ and *GsdmD*^*-/-*^ lung sections when compared to WT, with a statistically significant difference seen when comparing WT and *Casp11*^*-/-*^, but not between *Casp11*^*-/-*^ and *GsdmD*^*-/-*^ mice (**Fig. 3i**). These findings were corroborated by flow cytometric analysis quantifying the percentage of Ly6G^high^ neutrophils among the CD45^+^ immune cells in lung single cell suspensions from SAR-CoV2-infected WT, *Casp11*^*-/-*^ and *GsdmD*^*-/-*^ mice (**Fig. 3j,k**).

One of the main neutrophil-mediated functions is formation of Neutrophil extracellular traps (NETs) which are concentrated chromatin released to immobilize pathogens, and can trigger immunothrombosis especially during SARS-CoV-2 infection through platelet-neutrophil interactions^22^. To determine if CASP11 and GSDMD modulate neutrophil functions during SARS-CoV-2 infection, WT, *Casp11*^*-/-*^ and *Gsdmd*^*-/-*^ neutrophils were treated with PMA (**Fig. 4a**,), or culture supernatants of WT epithelial cells infected with SARS-CoV-2 (**Supplementary Fig. 3**). Notably, *Casp11*^*-/-*^ neutrophils failed to form NETs in response to all conditions. In contrast, *Gsdmd*^*-/-*^ and WT neutrophils formed NETs in response to all conditions (**Fig. 4a, Supplementary Fig. 3**). Together, our data demonstrate that lungs of SARS-CoV-2-infected *Casp11*^*-/-*^ mice contain fewer neutrophils than infected WT lungs, and additionally, that *Casp11*^*-/-*^ neutrophils largely fail to activate and undergo NETosis.

**Figure 4:**
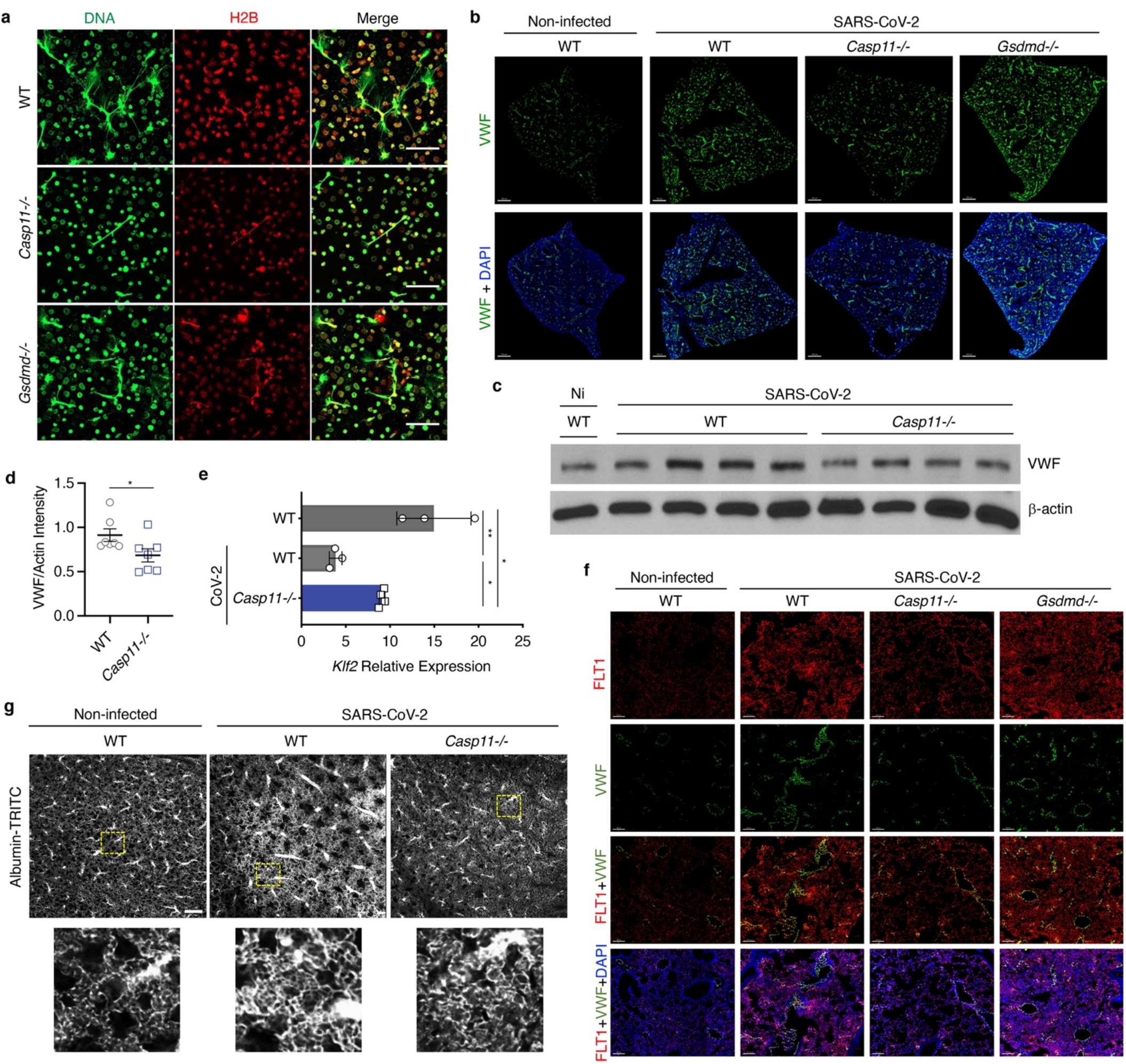
*Casp11*^*-/-*^ neutrophils undergo less NETosis and *Casp11*^*-/-*^ mice show decreased indicators of coagulopathy in lungs after SARS-CoV-2 infection. **a**, Neutrophils from WT, *Casp11*^-/-^ and *Gsdmd*^*-/-*^ mice were treated with PMA and NET formation was visualized by staining with anti-mouse Histone 2b (red), and anti-dsDNA (green). Images were captured at 60x magnification. **b-g**, WT, *Gsdmd*^*-/-*^, and *Casp11*^*-/-*^ mice were infected with SARS-CoV-2 (MA10, 10^5^ pfu). Lungs were collected at day 4 post-infection. **b**. RNA for *VWF* (green) was stained by RNAscope *in situ* hybridization, and nuclei are stained with DAPI (blue). Images were captured by a 20x objective in a 3D stitched panoramic view. Intensity projection images were created using IMARIS software. **c**, Western blotting of lung homogenates from non-infected WT and SARS-CoV-2-infected WT and *Casp11*^*-/-*^ mice as described in **a** were quantified in **d**, unpaired t test. **e**, qRT-PCR quantification of KLF2 in the lungs of mice as described in **b**, unpaired t test. **f**, Confocal microscopy for the colocalization of *VWF* RNA (green) with endothelial VEGF receptor subtype 1 (FLT1, red) in the lungs of mice as described in **b**. Nuclei were stained with DAPI (blue). Images were captured with a 20x objective in a z-stack 3D view and visualized using intensity projection function of IMARIS software. **g**, Vasculature imaging of intact lungs 4 days post-infection. Panel below, higher-magnification view of the regions in yellow boxes. Scale bar, 200 microns. *p<0.05, **p<0.005.

### The lack of CASP11 reduces von Willebrand factor levels and increases vascular integrity in response to SARS-CoV-2

SARS-CoV-2 infection is accompanied by long-term sequela mediated in part by vascular damage and thrombosis^23^. Given that we noted decreased neutrophil gene signatures in *Casp11*^*-/-*^ lungs upon infection, and since tissue infiltration by neutrophils can activate blood clotting cascades and thrombosis^24,25^, we examined whether the production of von Willebrand factor (VWF), which is essential to thrombus initiation and stabilization, is regulated by CASP11. Using RNAscope in situ hybridization (ISH) technology, we found significantly more blood vessels expressing VWF mRNA in the lung vascular architecture of SARS-CoV-2 infected WT mice when compared to *Casp11*^*-/-*^ lungs at day 4 post-infection (**Fig. 4b, Supplementary Fig. 4)**. Immunoblot analysis of lung homogenates confirmed that VWF was significantly lower in lungs of *Casp11*^-/-^ mice when compared to WT lungs (**Fig. 4c,d**). Notably, lung sections from SARS-CoV-2-infected *Gsdmd*^*-/-*^ showed more staining for VWF than *Casp11*^*-/-*^ mice (**Fig. 4b)**. Therefore, CASP11 is required for the accumulation of VWF in the lungs during SARS-CoV-2 infection. To determine the source of VWF, lung sections from *Casp11*^*-/-*^ and *Gsdmd*^*-/-*^ mice were processed for the simultaneous detection of endothelial marker VEGF receptor 1 (FLT1) and VWF mRNA^26^. We found that VWF RNA colocalized with FLT1, which was also upregulated in WT and *Gsdmd*^*-/-*^ but not *Casp11*^*-/-*^ lung sections (**Fig. 4f, Supplementary Fig. 5**). Furthermore, we examined the expression of Kruppel-Like Factor 2 (KLF2) in WT and *Casp11*^*-/-*^ SARS-CoV-2-infected lungs. KLF2 is an endothelial protective transcription factor that exerts anti-inflammatory and anti-thrombotic functions in the vascular endothelial cell and maintains the integrity of the endothelial vasculature. We found that KLF2 expression is significantly reduced after SARS-CoV-2-infection in WT lungs, but largely preserved in *Casp11*^*-/-*^ infected lungs **(Fig. 4e)**. Moreover, we examined the vascular architecture in the cleared lungs of SARS-CoV-2 infected mice by using fluorophore conjugated albumin and tissue clearing (**Supplementary Fig 6**). The vascular tracing revealed distinctive vascular features in WT SARS-CoV-2-infected lungs with pronounced vascular thickening and angiogenesis/neovascularization **(Fig. 4g)**. In stark contrast, *Casp11*^*-/-*^ infected lung vasculature did not show these abnormalities, confirming less endothelial damage/dysfunction **(Fig. 4g)**. Taken together, we conclude that CASP11 contributes to endothelial injury and instigation of the coagulation cascade during SARS-CoV-2 infection.

## Discussion

The medical and research communities have met challenges in identifying specific inflammatory mediators that can be targeted to ameliorate disease without impairing beneficial aspects of the immune response, such as viral clearance. A major impediment to mechanistic research in this regard has been the difficulty in infecting mouse models with SARS-CoV-2. Here we utilized the mouse-adapted SARS-CoV-2 (strain MA10)^15^ that was plaque purified, grown in Vero-TMPRSS2 cells, and sequenced to ensure that it lacks the attenuating tissue culture adaptations present in stocks of the virus grown in standard Vero cells, the most commonly used cell line for SARS-CoV-2 propagation^27^. Our extensive purification regimen allowed us to achieve measurable pathogenicity in C57BL/6 mice and to infect gene knockout (KO) animals for mechanistic research *in vivo*. This manuscript thus represents one of the first *in vivo* studies performed with SARS-CoV-2 in specific KO animals.

The active inflammasome complex has been implicated in many disease conditions and infections, including SARS-CoV-2^7,13^. Cell culture experiments identified a minor role for the canonical inflammasome member caspase-1 (CASP1) in SARS-CoV-2 infection^13^. On the other hand, CASP11, a member of the non-canonical inflammasome, has not been previously investigated in this context *in vitro* or *in vivo*. CASP11 is not expressed by resting cells, yet it is induced by bacterial infection and several cytokines^19,28,29^. We mined available clinical data and found that the expression of human CASP4 in COVID-19 testing swab material correlates with the severity of SARS-CoV-2 infection. Additionally, we found that the expression of CASP4 is elevated in lung sections of SARS-CoV-2 patients. Similarly, mouse CASP11 is upregulated in the lungs of WT mice in response to SARS-CoV-2. We previously reported that CASP11 restricts

*Legionella pneumophila* and *Burkholderia cenocepacia* infections by regulating actin dynamics^9^. CASP11 recognizes bacterial lipopolysaccharide (LPS) in the cytosol leading to downstream activation of CASP1 and IL-1β^30^. However, the role of CASP11 is not restricted to Gram-negative bacteria that produce LPS^9^, since we found that CASP11 is exploited by the Gram-positive bacteria methicillin-resistant *Staphylococcus aureus* (MRSA), to survive in macrophages^18^. In these cases, CASP11 regulates the functions of actin machinery to affect vesicular trafficking and cell migration. While it is possible that reduced neutrophil infiltration in SARS-CoV-2-infected *Casp11*^*-/-*^ lungs is due to reduced cytokine and chemokine levels in the lungs, we have also previously shown that even with exogenous addition of chemoattractants, *Casp11*^*-/-*^ immune cells, particularly neutrophils, fail to travel to the inflammation site due to an inherent defect in cell movement^19^. Our lung histology and flow cytometry data show that neutrophil reduction in *Casp11*^*-/-*^ and *Gsdmd*^*-/-*^ mice is comparable, yet the pathology in these animals is different. Our transcriptional profiling revealed a defect in cytokine responses, cellular recruitment, and immune activation in the absence of CASP11 demonstrating that *Casp11*^*-/-*^ neutrophils may be non-functional when compared to WT and *Gsdmd*^*-/-*^ neutrophils, a notion that is supported by the lack of NETosis in *Casp11*^*-/-*^ neutrophils. On the other hand, GSDMD, which is considered the best characterized effector of CASP11 and CASP4^12^, did not contribute to the lung pathology of SARS-CoV-2-infected mice explaining why clinical trials using GSDMD inhibitors were not successful^13^. Hence, our data suggest that CASP11 mediates many functions that are not executed by GSDMD.

The lungs of human patients infected with SARS-CoV-2 show diffuse immune cell infiltration, alveolar damage, alveolar edema and proteinaceous exudates and destruction of endothelial cells, indicative of acute respiratory distress syndrome (ARDS)^1,2^. Similar findings are detected in WT and *Gsdmd*^-/-^ mice while lung morphology appear healthier in *Casp11*^-/-^ mice after SARS-CoV-2 infection. In addition, there is less weight loss, with fast recovery to normal weight in *Casp-11*^-/-^ mice, when compared with WT and *Gsdmd*^-/-^ mice, which are slower to recover. Importantly, the differences in disease severity are not due to changes in viral burden among different genotypes. This is consistent with a lack of changes in global ISG expression in WT versus *Casp11*^-/-^ or *Gsdmd*^*-/-*^ lungs, which are genes implicated in viral resistance and clearance. Instead, we observed reduced inflammation and lung pathology dependent on CASP11 irrespective of viral loads. In *Casp11*^*-/-*^, but not *Gsdmd*^*-/-*^ SARS-CoV-2-infected mice, cytokines including *Cxcl1, Cxcl2, Cxcl14*, which are involved in neutrophil and monocyte recruitment^31^, were significantly down-regulated. However, there was no significant difference in expression of IL-1β between *Casp11*^*-/-*^ and *Gsdmd*^*-/-*^ mice. *In vitro*, IL-1β was barely detectable in the supernatants of macrophages infected with SARS-CoV-2. This is explained by a recent publication demonstrating that SARS-CoV-2 nucleocapsid inhibits the cleavage of GSDMD in infected cells and hence prevents the release of IL-1β^32^. In addition, our data demonstrate that IL-6 is elevated in infected lungs in a CASP11-dependent manner. IL-6 was identified during COVID-19 pandemic as being a highly upregulated mediator of disease severity in ill patients. Moreover, high levels of IL-6 can also activate the coagulation system and increase vascular permeability^3^.

Post-mortem studies have highlighted disseminated micro-thrombi which together with increased mortality, morbidity and long-term sequel from SARS-CoV-2 infection, are considered hallmarks of severe COVID-19^33,34^. Currently, the administration of an anticoagulant such as heparin for all hospitalized COVID-19 patients is associated with lower mortality rates and better prognosis^3^. Typically, endothelial activation and damage leads to increased VWF production and this activates the coagulation cascade, along with extensive NETosis elicited by neutrophils leading to prothrombotic events^23,35^. Importantly, we found here that lungs from *Casp11*^*-/-*^ mice accumulate significantly less VWF in response to SARS-CoV-2 infection, which is largely confined to what appears to be lining of blood vessels. In contrast, the distribution of VWF in WT lungs was intense and diffuse suggesting the presence of vascular damage. Notably, lung sections from *Gsdmd*^*-/-*^ expressed more VWF than those from *Casp11*^*-/-*^ mice. To further evaluate endothelial damage, we determined the expression of the transcription factor KLF2. Recent reports have linked the vascular injury that is associated with SARS-CoV-2 to the reduction in the expression of KLF2 in lung endothelial cells^36^. We found that KLF2 levels are largely preserved in the *Casp11*^*-/-*^ lungs but are significantly reduced in WT and *Gsdmd*^*-/-*^ lungs. Moreover, the vascular abnormalities we detected on lung vascular tracing indicate severe endothelial damage and endothelialitis in WT SARS-CoV-2-infected lungs. These vascular features resemble the intussusceptive angiogenesis that has been described in SARS-CoV-2-infected human lungs^37,38^. Importantly, the inhibition of angiogenesis through targeting vascular endothelial growth factor (VEGF) has been proven beneficial in patients with severe SARS-CoV-2^39^. Notably, we have found less expression of VEGF receptor 1 (FLT1) with less angiogenesis and neovascularization in the infected *Casp11*^*-/-*^ lungs compared to WT and *Gsdmd*^*-/-*^ lungs. Our data demonstrate a previously unrecognized function for CASP11 which is the promotion of coagulation pathways and endothelial dysfunction that lead to thrombotic events.

Together, our findings suggest that targeting the CASP11 homologue, human CASP4, during COVID-19 will prevent severe pneumonia, inflammation, tissue damage as well as thrombosis and accompanying repercussions such as low oxygen, lung failure, need for ventilators and perhaps long-term sequela. These advantageous effects will be achieved without compromising viral clearance. It is also plausible that the level of expression of CASP4 could serve as a biomarker to identify patients who will succumb to severe Covid. Targeting CASP4 alone can achieve benefits that will exceed and replace the administration of a large number of individual anti-inflammatory agents and anti-thrombotics given to SARS-CoV-2 patients. Further research is needed to develop therapeutics in this regard.

## Materials and Methods

### Biosafety

All experiments with live SARS-CoV-2 were performed in the OSU BSL3 biocontainment facility. All procedures were approved by the OSU BSL3 Operations/Advisory Group, the OSU Institutional Biosafety Officer, and the OSU Institutional Biosafety Committee.

### Viruses and titers

Mouse adapted SARS-CoV-2, variant strain MA10^15^, generated by the laboratory of Dr. Ralph Baric (University of North Carolina) was provided by BEI Resources (Cat # NR-55329). SARS-CoV-2 strain USA-WA1/2020 was also provided by BEI Resources (Cat # NR-52281). Viral stocks from BEI Resources were plaque purified on Vero E6 cells to identify plaques lacking mutations in the polybasic cleavage site of the Spike protein via sequencing. Non-mutated clones were propagated on Vero E6 cells stably expressing TMPRSS2 (provided by Dr. Shan-Lu Liu, The Ohio State University). Virus aliquots were flash frozen in liquid nitrogen and stored at -80 C. Virus stocks were sequenced to confirm a lack of tissue culture adaptation in the polybasic cleavage site. Virus stocks and tissue homogenates were titered on Vero E6 cells.

### Mice

C57BL/6 wild-type (WT) mice were obtained from the Jackson Laboratory (Bar Harbor, ME, USA). *Casp11*^-/-^ mice were generously given by Dr. Yuan at Harvard Medical School, Boston, MA, USA^106^. *Gsdmd*^-/-^ mice were a gift from Dr. Thirumala-Devi Kanneganti at St. Jude Children’s Research Hospital, Memphis, TN, USA. K18-hACE2 mice^40^ were purchased from Jackson Laboratories. All infections were performed intranasally on anesthetized mice with viruses diluted in sterile saline. All mice were housed in a pathogen-free facility, and experiments were conducted with approval from the Animal Care and Use Committee at the Ohio State University (Columbus, OH, USA) which is accredited by AAALAC International according to guidelines of the Public Health Service as issued in the Guide for the Care and Use of Laboratory Animals.

### Derivation of single cell suspension and primary lung macrophages

Lungs were perfused with cold PBS to remove circulating intravascular WBCs. Lungs were dissected into single lobes before being dissociated into single cell suspension using gentleMACS octo-dissociator and Miltenyi lung dissociation kit (Miltenyi Biotec, 130-095-927). Red blood cells (RBCs) were lysed by incubating cells in 2 ml ACK buffer for 5 min at room temperature. After RBCs lysis, cells were washed in DPBS containing 1% BSA. The single cell suspension was centrifuged, and the cell pellets were washed twice with PBS. Cell pellets were further suspended in 0.5 ml of PBS 1%BSA. This was followed by CD11b magnetic bead (Miltenyi Biotec, 130-049-601) isolation technique to positively select for macrophage expressing the pan-macrophage/monocyte CD11b marker.

### Flow cytometry

Single cell suspension from the previous step was stained with fluorophore conjugated antibodies for fluorometric analysis as described before ^41^.

### Murine tracheobronchial epithelial 3D cultures

Murine trachea and bronchioles were dissected from two mice each of C57Bl/6 WT, *Casp11*^*-/-*^ and *Gsdmd*^*-/-*^. Isolation of tracheobronchial epithelial cells was as follows. Tissues were washed, and tracheas were incubated overnight in Ham’s F12, 1% penicillin/streptomycin, 1% amphotericin B (Fisher Scientific, #15290018) and Pronase from Streptomyces griseus (Sigma Aldrich, #10165921001) solution. Digestion of trachea and bronchioles were neutralized with 10% fetal bovine solution (FBS; Life Technologies, #10438026) and tracheal airway cells were gently scraped. Cells were washed three times in Ham’s F12, 10% FBS and 1% penicillin/streptomycin solution and further digested in DNase I solution (Sigma Aldrich, #DN25-10) in Ham’s F12 with 10mg/mL bovine serum albumin (Fisher Scientific, #BP9706). Airway cells were then washed with Murine Tracheobronchial Epithelial Cell (MTEC) base medium [1:1 Ham’s F12: DMEM (Fisher Scientific, #11995065), plus 10% FBS, 1% penicillin/streptomycin, 50μg/mL gentamicin (Life Technologies, #15710064), and 0.03% w/v NaHCO3]. Cells were plated in a T25 flask (Fisher Scientific, #1012610) overnight in MTEC medium at 37oC, 5% CO2. The next day, medium was switched to 1:1 of MTEC and PneumaCult-Ex PLUS medium (StemCell Technologies, #05040) and fed every other day until expansion of cells to ∼80% confluent. Epithelial cells were then trypsinized twice with TrypLE Express (ThermoFisher, #12605010) to remove residual fibroblast cells and seeded at a density of 50,000 cells per transwell in Corning 6.5mm 24-well transwells (Fisher Scientific, #07200154) in 1:1 MTEC:PneumaCult-Ex PLUS medium. Cells were fed for 4-5 days until airlifted and continued to be grown at air-liquid interface (ALI) with PneumaCult ALI medium (StemCell Technologies, #05001) until fully differentiated (4 weeks)

### Immunoblotting

Protein extraction from lung tissue was performed using TRIzol reagent (Thermo Fisher Scientific, 15596026) according to the manufacturer’s instructions. Equal amounts of protein were separated by SDS-PAGE and transferred to a polyvinylidene fluoride (PVDF) membrane. Membranes were incubated overnight with antibodies against CASP11 (Cell Signaling Technology, 14340), VWF (Protein tech, 11778-1-AP), and β-Actin (Cell Signaling Technology,3700). Corresponding secondary antibodies conjugated with horseradish peroxidase in combination with enhanced chemiluminescence reagent (Amersham, RPN2209,) were used to visualize protein bands. Densitometry analyses were performed by normalizing target protein bands to their respective loading control (β-Actin) using ImageJ software as previously described ^19,42^.

### ELISAs

Cytokine/chemoking ELISAs were performed on lung homogenates or macrophage supernatants using R&D Systems Duoset ELISA kits (IL-6, DY406; IL-1b, DY401; CXCL1, DY453) according to the manufacturer’s instructions.

### Histology

Lungs were removed from infected mice, and fixed in 10% formalin at room temperature. Sample preparation, processing, hematoxylin and eosin staining (H&E), and semi-quantitative slide evaluation using ordinal grading scales was performed as previously described^59^. Lungs used for immunofluorescence staining and RNAscope^®^ ISH technique were embedded in OCT and flash frozen while lung tissue used for IHC was embedded in paraffin blocks.

### Immunohistochemistry (IHC) and Immunofluorescence (IF) staining for mouse tissue

Immunofluorescence (IF) staining of mouse lung sections has been performed as previously described^42^. Slides were washed 3 times for 15 min with PBS to remove residual OCT. The sections were then incubated in the blocking solution (PBS containing 10% donkey serum (cat no: S30-100ml, Millipore Sigma), 2% BSA (cat no: BP1600-100, Fisher Scientific) and 0.3% Triton X-100 (cat no: BP151-100, Fisher Scientific) for 2 h at room temperature. Sections were then transferred to blocking solution containing the primary antibody against IL1β (GeneTex, GTX74034), and incubated overnight at 4ºC. After that, sections were washed with PBS 3X for 15 min each. Then, they were incubated with the blocking solution containing the secondary antibody, for 2 h at room temperature. DAPI (cat no: D1306, Fisher Scientific) was added to the staining solution in the last 15 min of incubation at a final concentration (5ug/ml). Finally, sections were washed with PBS 3X for 15 min. Antifade mounting media (cat no: P36934, Thermo Fisher Scientific) was added before cover-slipped. For IHC, Ly6G (Abcam, ab25377) and SARS-CoV-2 nucleocapsid protein (GeneTex, GTX635686) primary antibodies were used. All the stainings were performed at Histowiz, Inc Brooklyn, using the Leica Bond RX automated stainer (Leica Microsystems). The slides were dewaxed using xylene and alcohol based dewaxing solutions. Epitope retrieval was performed by heat-induced epitope retrieval (HIER) of the formalin-fixed, paraffin-embedded tissue using citrate based pH 6 solution for 40 mins at 95 C. The tissues were first incubated with peroxide block buffer (Leica Microsystems), followed by incubation with the rabbit Caspase 4 antibody (Novus Bio NBP1-87681) at 1:700 dilution for 30mins, followed by DAB rabbit secondary reagents: polymer, DAB refine and hematoxylin (Leica Microsystems). The slides were dried, coverslipped and visualized using a Leica Aperio AT2 slide scanner (Leica Microsystems).”

### RNAscope In situ hybridization (ISH)

Lung tissue was fixed and embedded in OCT as described above. Sections of 15 microns thickness were mounted on Plus charged slides. ISH was performed using RNAscope Multiplex Fluorescent Reagent Kit v2 (Advanced Cell Diagnostics, Cat. No. 323100) as described before^43^. All incubations between 40 and 60° C were conducted using an ACD HybEZ II Hybridization System with an EZ-Batch Slide System (Advanced Cell Diagnostics; cat# 321710). Slides were washed in PBS twice to remove any residual OCT then baked at 60°C for 30 minutes. Baked slides were subsequently post fixed in cold 10% formalin for 15 minutes then washed and treated with Hydrogen Peroxide solution (10 min at RT; Advanced Cell Diagnostics, cat# 322335). After being rinsed twice with ddH2O, sections were incubated in RNAscope Target Retrieval Solution (98 C for 5 min; Advanced Cell Diagnostics, cat# 322001) and rinsed 3 times. Next, a hydrophobic barrier was created around the tissue using an ImmEdge Pen (Advanced Cell Diagnostics; cat# 310018), and slides were incubated with RNAscope® Protease III (30 min at 40 C; Advanced Cell Diagnostics, cat# 322337), and subsequently incubated with RNAscope target probes VWF(cat# 499111), FLT1(cat# 415541-C2), Casp4/Casp11 (cat# 589511) for 2 hours at 40°. Next, slides were washed twice with 1X Wash Buffer (Advanced Cell Diagnostics, cat# 310091; 2 min/rinse at RT) followed by sequential tissue application of the following: RNAscope Multiplex FL v2 Amp 1 (Advanced Cell Diagnostics, cat# 323101), RNAscope Multiplex FL v2 Amp 2 (Advanced Cell Diagnostics, cat# 323102), and RNAscope Multiplex FL v2 Amp 3 (Advanced Cell Diagnostics, cat# 323103). This was followed by application of RNAscope Multiplex FL v2 HRP C1or C2 (15 min at 40 C; Advanced Cell Diagnostics, cat#323104). Finally, Opal dyes (Opal 520 and 570 were used, Fisher Scientific; cat# NC1601877 and cat#NC601878) was then applied, 520 (Fisher Scientific; cat#NC1601877) diluted in RNAscope TSA buffer (Advanced Cell Diagnostics, cat# 322809) for 30 min at 40 C. HRP blocker was subsequently added to halt the reaction. Finally, slides were incubated with DAPI, coverslipped with ProLong Gold Antifade Mountant (Fisher Scientific, cat# P36930), and stored at 4 C until image acquisition.

### Confocal imaging and analysis

Fluorescent images were captured on Olympus FV 3000 inverted microscope with a motorized stage. A 2x objective was used to create a map of the lung section in the X,Y dimension. This was followed by using 20x objective to create a stitched z stacked three dimensional panoramic view of the lung section. Images were taken by using the 488 nm, 543 nm, and 405 nm (for DAPI) lasers. Image reconstructions of z-stacks and intensity projection images (IPI) were generated in Imaris software (Bitplane, Inc.). *Flt1* mRNA expression was quantified using spot function in IMARIS. Number of cells was also quantified via the spot functions.

### Vasculature labeling with conjugated albumin

The mouse vasculature was labeled as reported by Di Giovanna *et al*.^44^. Briefly, mice were transcardially perfused with 10% formalin in phosphate buffered saline (PBS). Mice were then perfused with 5ml of 0.05% albumin-tetramethylrhodamine isothiocyanate bovine (A2289, Sigma) in 2% gelatin from porcine skin (G1890, Sigma). At the time of injection, the temperature of the gel solution was kept at 45°C. After clamping the heart, mice were placed on ice to lower the body temperature and allow gel formation. Lungs were post-fixed in 10% formalin for 10 days. The unsectioned lungs were then cleared using the advanced CUBIC protocol^45^ and imaged using a confocal microscope (C2, Nikon).

### NET Formation Assay

Bone marrow was collected from WT, *Gsdmd*^*-/-*^ or *Casp11*^*-/-*^ mice, then neutrophils were negatively selected by using the EasySep™ mouse neutrophil enrichment kit (STEM cell technologies, #19762A), and 200,000 neutrophils/well were plated in 24-well plate on fibronectin coated glass coverslip. Polymorphonuclear neutrophils (PMNs) were stimulated for 4 h with 100 nM PMA (Sigma-Aldrich, #P8139-10MG) or conditioned media from SARS-CoV-2-infected epithelial cells. The cells were fixed with 4% paraformaldehyde, permeabilized with 0.2% of Triton X-100 for 10 minutes and blocked with 10% goat serum for 30 min at RT. For the visualization of Neutrophils Extracellular Traps (NETs), neutrophils were stained with rabbit anti-mouse Histone 2b (Abcam, #ab1790), mouse anti-dsDNA (Abcam, #ab27156), goat anti-rabbit IgG Alexa Fluor 555 (Thermofisher, #A32732), goat anti-mouse IgG Alexa Fluor 488 (Abcam, #ab150113) and wheat germ agglutinin (WGA) Alexa Fluor 350 (Thermofisher, #W11263). The coverslips were mounted with Fluoroshield Mounting Medium (Abcam, #ab104135). The cells were visualized by confocal microscopy (Zeiss 800 Confocal microscope).

### RNAseq and data analysis

Total RNA was extracted from day 2 SARS-CoV-2 WT, *Casp11*^-/-^, and *Gsdmd*^-/-^ infected lungs by TRIzol reagent (Thermo Fisher Scientific, 15596026) according to the manufacturer’s instructions. RNA cleaning and concentration was done using Zymo Research, RNA Clean & Concentrator™-5 kit (cat# R1015) following the manufacturer’s protocol. Fluorometric quantification of RNA and RNA integrity analysis was carried out using RNA Biochip and Qubit RNA Fluorescence Dye (Invitrogen). cDNA libraries were generated using NEBNext® Ultra™ II Directional (stranded) RNA Library Prep Kit for Illumina (NEB #E7760L). Ribosomal RNA was removed using NEBNext rRNA Depletion Kit (human, mouse, rat) (E #E6310X). Libraries were indexed using NEBNext Multiplex Oligos for Illumina Unique Dual Index Primer Pairs (NEB #644OS/L). Library prep generated cDNA was quantified and analyzed using Agilent DNA chip and Qubit DNA dye. Ribo-depleted total transcriptome libraries were sequenced on an Illumina NovaSeq SP flow cell (paired-end 150bp format; 35-40 million clusters, equivalent to 70-80 million reads. Library preparation, QC, and sequencing was carried out at Nationwide Children’s Hospital genomic core.

Sequencing data processing and analysis was performed by the Bioinformatics Shared Resource Group (BISR) at the Ohio State University using previously published pipelines ^46^. Briefly, raw RNAseq data (fastq) were aligned to mouse reference genome (GRCh38) using hisat2 (v2.1.0) ^47^ and converted to counts using the ‘subread’ package (v1.5.1) ^48^ in R. In the case of multimapped reads, the primary alignment was counted. Low expressed counts were excluded if more than half of the samples did not meet the inclusion criteria (2 CPM). Data were normalized using ‘voom’ and statistical analysis for differential expression was performed with ‘limma’ ^49^. For data visualization, DESeq2 rlog transformation was used for principal component analysis (PCA). Volcano plots were generated with ‘EnhancedVolcano’ and heatmaps were generated ‘ComplexHeatmap’ using R. Functional enrichment performed with Ingenuity Pathway Analysis (Qiagen) to enrich for IPA Canonical Pathways, ‘clusterProfiler’ to generate enrichment maps ^46^, and EnrichR ^50^.

## Statistical analysis

Data were analyzed using GraphPad Prism 8.3.0. All figures display mean and standard deviation (SD) or standard error of the mean (SEM) from independent experiments as indicated in the figure legends. Comparisons between groups were conducted with either upaired t-test or ANOVA followed by Tukey’s multiple comparisons test. Adjusted P<0.05 was considered statistically significant.

## Data availability

Data shared through Gene Expression Omnibus with accession number GSE184678.

## Author Contributions

Conceptualization, A.O.A., J.S.Y.; Experiments and data acquisition, M.E., A.Z., A.D.K., S.E., A.B., E.A., E.K., C.C., K.D., O.W., J.K., A.E., P.D. E.K, E.A.H., E.C.-B., P.B.; Generation of critical reagents and patient samples, M.K.C, M.L., J.L., M.P., J.Z., J.Q., A.T.; Data Analysis, M.E., A.Z., M.P., A.W., A.F., A.O.A, J.S.Y. ; Writing – Original draft, A.O.A, J.S.Y., A.F.; Writing – Review and editing, all authors.; Project Administration, A.O.A., J.S.Y.; Supervision, A.O.A., J.S.Y, E.C.-B., P.B.; Funding Acquisition, A.O.A and J.S.Y.

## Acknowledgments

Studies in the Amer laboratory are supported by AI24121, HL127651, NIH Covid supplement and a Pilot Grant from the OSU Department of Microbial Infection and Immunity. A.B. is supported in part by the C3 training grant. Studies in the Yount laboratory are supported by NIAID grants AI130110, AI151230, AI142256, AI146690, and AI151230, as well as by the American Lung Association’s COVID-19 and Emerging Respiratory Viruses Research Award. Ashley Zani is supported by an NSF-GRFP fellowship. The authors declare that they have no conflict of interests.

## Supplementary figures

**Supplementary Fig. 1:**
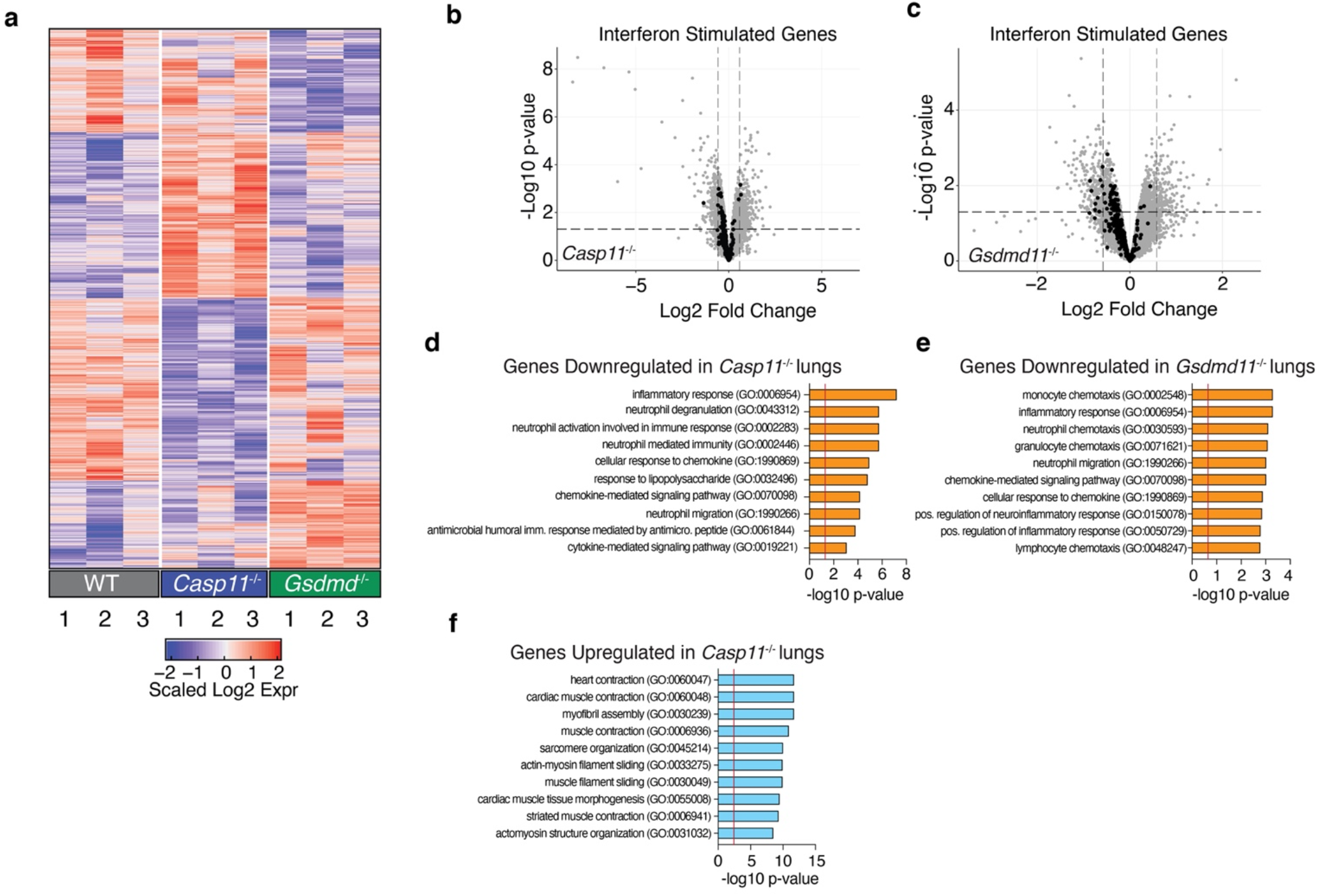
Changes in inflammatory responses in *Casp11*^*-/-*^ and *Gsdmd*^*-/-*^ SARS-CoV-2-infected lungs. **a**, Heat map of significant gene expression changes (p-value <0.05). Depicted genes were chosen based on comparisons relative to WT. Color indicates relative upregulation (red) or downregulation (blue) in gene expression. **b-c**, Statistical analysis of ISG expression in *Casp11*^-/-^ and *Gsdmd*^*-/-*^ infected lungs relative to WT. Each point represents transcripts within the dataset. 300 IFNb-responsive ISGs are highlighted in black. Dashed lines represent LFC and p-value cutoffs (LFC |0.58| and p-value 0.05). **d**, Functional enrichment analysis of the top 236 downregulated genes in *Casp11*^*-/-*^ SARS-CoV-2-infected lungs relative to infected WT. Red vertical line represents threshold of significance p-value 0.05. **e**, Functional enrichment analysis of the top 224 downregulated genes in *Gsdmd*^-/-^ infected lungs relative to WT infection. Red verical line represents threshold of significance (p-value <0.05). **f**, Functional enrichment analysis of 328 upregulated genes *in Casp11*^*-/-*^ infected lungs relative to WT infection.

**Supplementary Fig. 2:**
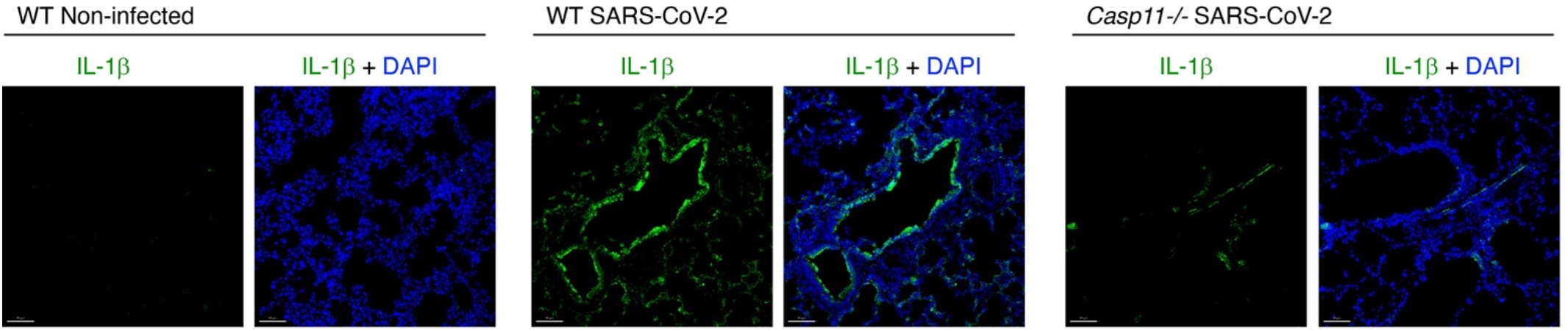
IL-1b production during SARS-CoV-2 infection is decreased in the absence of *Casp11*. WT and *Casp11*^*-/-*^ mice were infected with SARS-CoV-2 (MA10, 10^5^ pfu). Lungs were collected at day 4 post-infection. Lung tissue was sectioned and stained for IL-1β (green), and DAPI (blue).

**Supplementary Fig. 3:**
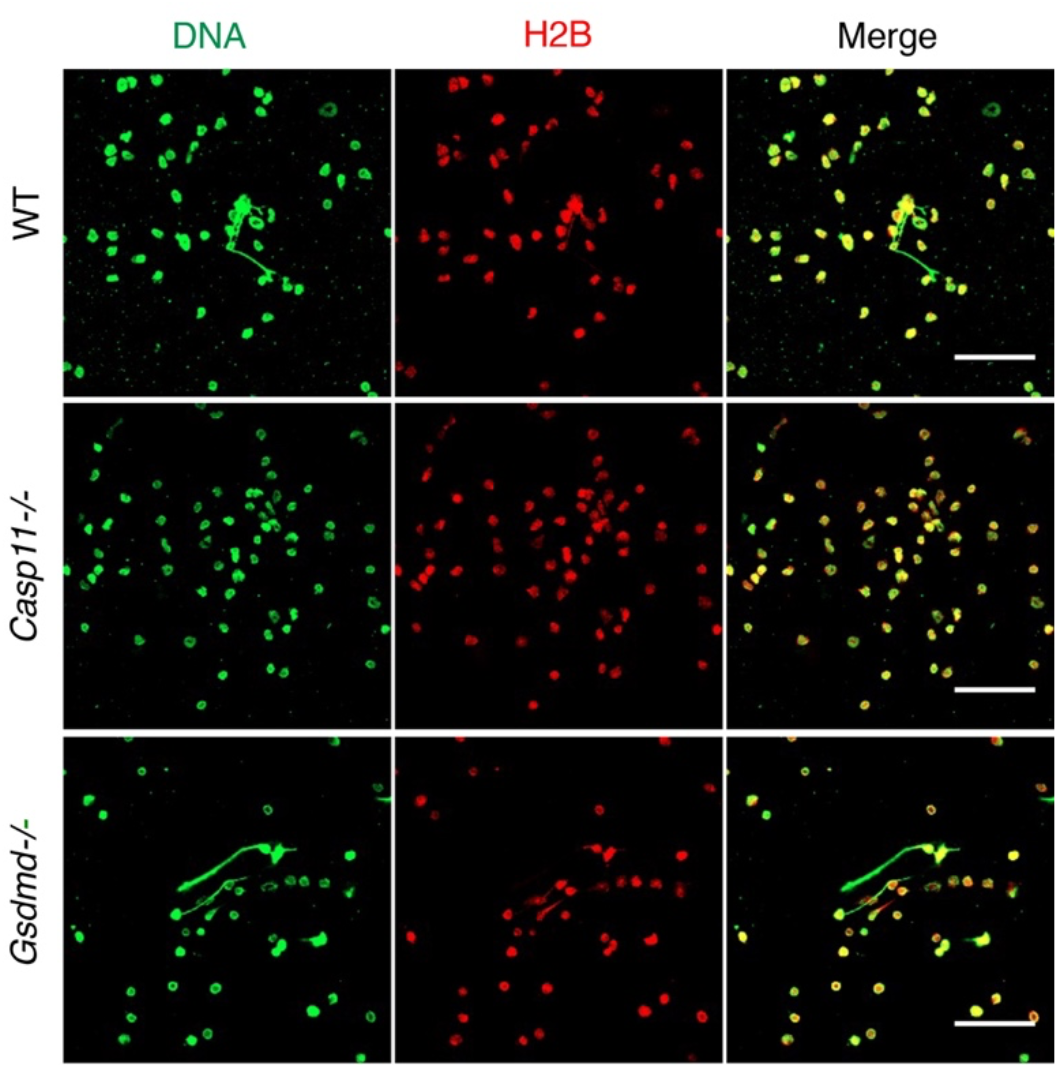
*Casp11*^*-/-*^ neutrophils are impaired in NET formation. Neutrophils from WT, *Casp11*^-/-^ and *Gsdmd*^*-/-*^ mice were treated with supernatants of SARS-CoV-2-infected epithelial cells from WT mice and NET formation was visualized with staining with anti-mouse Histone 2b (red) and anti-dsDNA (green). Images were captured at 60x magnification.

**Supplementary Fig. 4:**
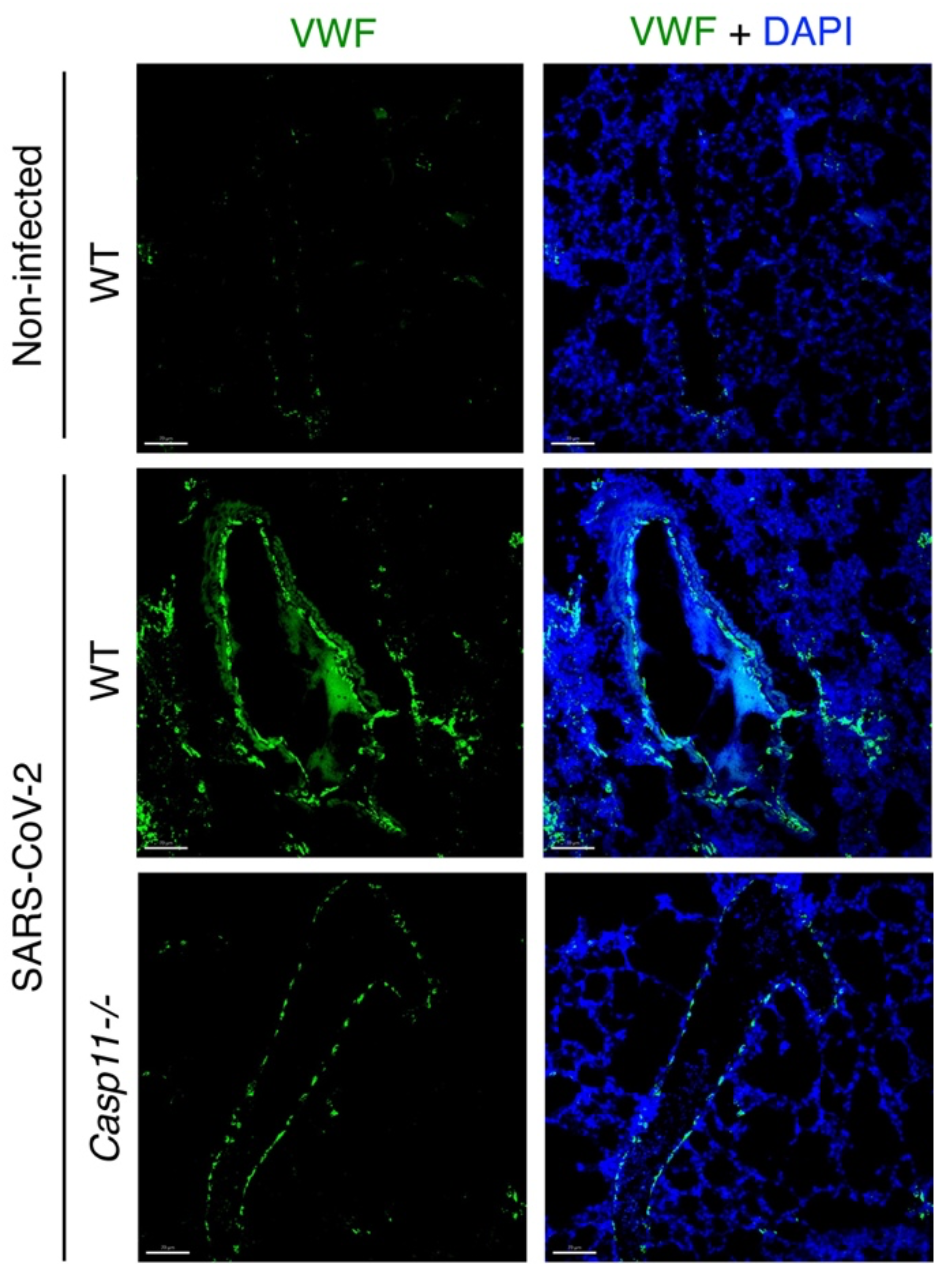
VWF accumulation at blood vessels during SARS-CoV-2 infection is decreased in the absence of *Casp11*. Mice were infected with SARS-CoV-2 (MA10, 10^5^ pfu). Lungs were collected at day 4 post-infection. RNA of *VWF* was detected by RNAscope *in situ* hybridization (green) and nuclei were staining with DAPI (blue). Images were captured by a 20x objective. Full lung stitched images are shown in main text **Fig. 4b**.

**Supplementary Fig. 5.**
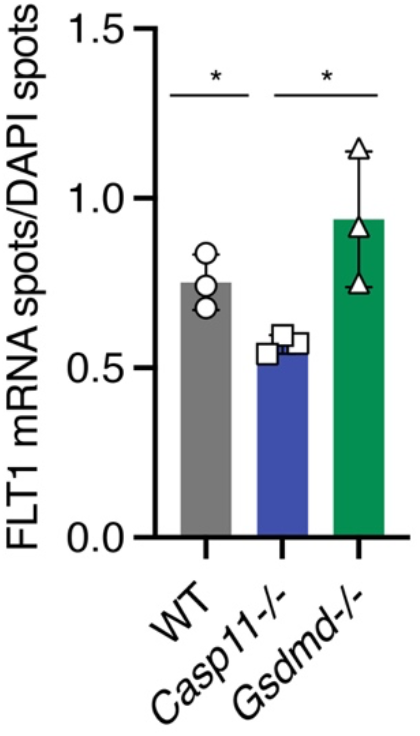
FLT1 is downregulated in *Casp11*^*-/-*^ SARS-CoV-2-infected lungs. Quantification of *in situ* hybridization RNAscope staining of endothelial VEGF receptor subtype 1 (FLT1) in lung sections. Mice were infected with SARS-CoV-2 (MA10, 10^5^ pfu). Lungs were collected at day 4 post-infection. Original Images were captured by a 20x objective in a 3D stitched panoramic view representing the whole lung in x,y and z in lung sections of samples described in **4f**. DAPI and FLT1 mRNA spots were quantified by using the spot function in IMARIS software. Unpaired t test. *p<0.05

**Supplementary Fig. 6:**
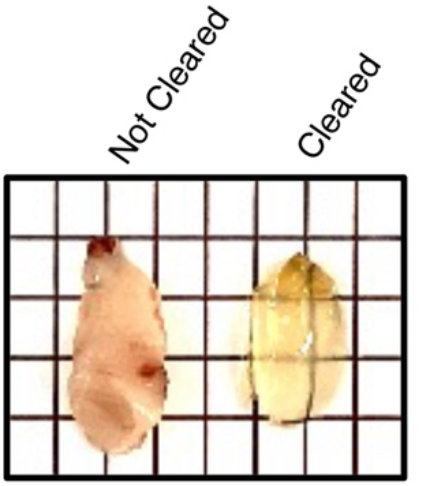
Example clearing of lungs for vascular imaging. Representative photograph of lungs with and without tissue clearing.

**Supplementary Fig. 7:**
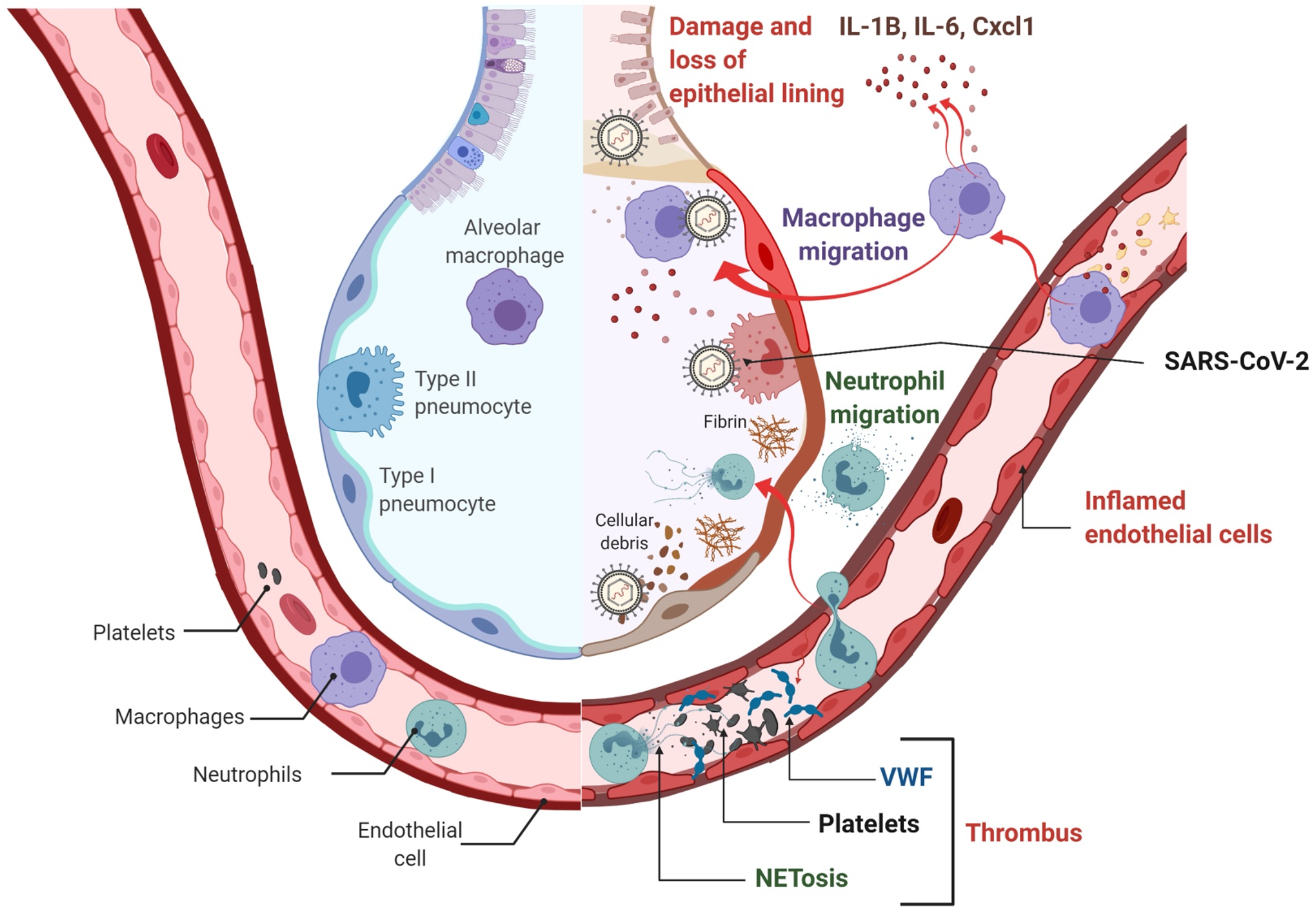
Casp11-mediates hyperinflammation, neutrophil infiltration, NETosis, thrombus formation and vascular damage during SARS-CoV-2 infection.

